# 1/*f* Neural Noise and Electrophysiological Indices of Contextual Prediction in Normative Aging

**DOI:** 10.1101/147058

**Authors:** S. Dave, T.A. Brothers, T.Y. Swaab

## Abstract

Prediction during language comprehension has increasingly been suggested to play a substantive role in efficient language processing. Emerging models have postulated that predictive mechanisms are enhanced when neural networks fire synchronously, but to date, this relationship has been investigated primarily through oscillatory activity in narrow frequency bands. A recently-developed measure proposed to reflect broadband neural activity – and thereby synchronous neuronal firing – is 1/*f* neural noise extracted from EEG spectral power. Previous research (Voytek et al., 2015) has indicated that this measure of 1/*f* neural noise changes across the lifespan, and these neural changes predict age-related behavioral impairments in visual working memory. Using a cross-sectional sample of young and older adults, we examined age-related changes in 1/*f* neural noise and whether this measure would predict ERP correlates of successful lexical prediction during discourse comprehension. 1/*f* neural noise across two different language tasks revealed high within-subject correlations, indicating that this measure can provide a reliable index of individualized patterns of neural activation. In addition to age, 1/*f* noise was a significant predictor of N400 effects of successful lexical prediction, but noise did not mediate age-related declines in other ERP effects. We discuss broader implications of these findings for theories of predictive processing, as well as potential applications of 1/*f* noise across research populations.

## Introduction

Studies of the effects of predictability during language comprehension in aging populations have traditionally examined cross-sectional differences that emerge when comparing groups of young versus older adult cohorts. Recent literature has sought to bolster this classic approach by considering the role of within-group variability. Across these studies, researchers have shown that neural indices of predictive processing vary as a function of age and maintenance of cognitive capacity (e.g., Dave, Brothers, Long, & Swaab, submitted; Delong, Groppe, Urbach, & Kutas, 2012; Federmeier, Kutas & Schul, 2010; Federmeier, McLennan, Ochoa, & Kutas, 2002; Huettig & Janse, 2016). Considerably less research has examined how maintenance of neural function impacts prediction across the lifespan, though a number of researchers have proposed that strong, coherent neuronal networks likely underlie preservation of predictive processing in older adults (Federmeier et al., 2010; Peelle, Troiani, Wingfield, & Grossman, 2010; Wlotko, Federmeier, & Kutas, 2012). Testing this hypothesis directly requires finding a reliable measure of neuronal network dynamics that can be correlated with both age and predictive processing during language comprehension. One potential measure has recently been proposed by Voytek and colleagues (2015): 1/*f* neural noise extracted from spectral power in the EEG signal. Voytek et al. found that 1/*f* neural noise positively correlated with age, and these neural changes explained age-related decline in visual working memory performance. Here we investigate whether there are similar age-related differences in 1/*f* neural noise during language comprehension, and if this measure is reliable on the individual level when collected during different tasks. Then we examine whether 1/*f* noise mediates the effect of age on neural indices of accurate prediction of words during discourse comprehension, i.e., lexical prediction.

### The Neural Noise Hypothesis of Aging

Neural noise has been defined as random, background electrical fluctuations within the central nervous system (Hong & Rebec, 2012; Li, Lindenberger, & Sikström, 2001; Serletis et al., 2011). Noise can be considered by comparison with the strength of the signal for relevant, or actively processed information; a nervous system is noisy if the ratio of signal strength to random background noise is low (signal detection theory in Nevin, 1969; reviewed in Swets, 2014). Noisy systems may emerge when connections between neurons are weakened or diffuse as a function of reduced arborization, neuronal loss, or inconsistent inhibition (Cremer & Zeef, 1987). The brains of older adults often show these types of losses and changes across neural connections (e.g., Cabeza, Nyberg, & Park, 2016; Fjell & Walhovd, 2010; Raz et al., 2005), underlying a long-held hypothesis (Crossman & Szafram, 1956) positing that the effective signal-to-noise ratio in neuronal networks decreases with age.

For decades, reaction times to degraded stimuli were the primary means by which researchers quantified neural noise (e.g., Creemer & Zeef, 1987; Salthouse & Lichty, 1985; Welford, 1981). However, behavioral measures may not accurately address connections in neural networks, and may only reflect age-related degradation in peripheral sensory systems (i.e, as opposed to cortical neural noise). An alternate method may be to measure neural noise directly and non-invasively using the electroencephalogram (EEG). Oscillatory neural activity recorded at the scalp is thought to arise from dynamic communication across ensembles of neurons, and a number of researchers have proposed that these oscillations measure synchrony in neuronal firing (e.g., Fries et al., 2007; Miller et al., 2014; Voytek & Knight, 2015). Recent studies have further indicated that population-level neuronal spiking correlates with oscillatory activity due to generalized broadband activation, as opposed to oscillations specific to any particular frequency band (e.g., Kreiman et al., 2006; Manning, Jacobs, Fried, & Kahana, 2009; Miller et al., 2007). In other words, neural noise may be reflected in the distribution of neuronal activation (or the power spectral density, PSD) across the frequency spectrum.

In order to test this hypothesis, Voytek et al. (2015) used EEG to investigate PSD in young and older adults. If, as suggested by the neural noise hypothesis, older adults have less synchronized or less simultaneous neuronal spiking relative to younger adults, this should be observed in the distribution of spectral power across frequencies. The characteristic distribution of EEG data in the frequency domain is inversely proportional (He, 2004; Miller, 2010; Voytek et al., 2015), such that a linear relationship can be approximated by calculating the log of the PSD. Voytek and colleagues showed that the slope of this linear relationship can serve as a measure of 1/*f neural noise*^1^, with more power at low frequencies relative to higher frequencies. In this study, older adults showed a flatter, less negative 1/*f* slope than younger adults. As PSD slopes are predicted to flatten when spiking activity is decoupled from oscillatory synchronization (Freeman & Zhai, 2009), these differences in 1/*f* neural noise have been interpreted as reflecting age-related declines in population-level synchrony across neuronal networks (Hong & Rebec, 2012; Podvalny et al., 2015; Voytek & Knight, 2015).

Voytek and colleagues (2015) found 1/*f* neural noise was higher for older adults performing a visual working memory task. In the present study, we examined whether similar age differences in 1/*f* noise could be observed during two language comprehension tasks. In addition to examining the effect of age on 1/*f* noise, we compared 1/*f* noise levels between two language tasks within individuals. In the first task (Comprehension Paradigm), participants were instructed to read single sentences for comprehension, while in the second task (Prediction Paradigm), participants were asked to try to predict the last word of a two-sentence passage. If 1/*f* noise represents a robust and reliable individual difference measure, we should observe strong correlations within subjects independent of differences in cognitive demands.

Finally, as Voytek et al. examined effects of 1/*f* neural noise on behavioral accuracy in their working memory task, this study tests if 1/*f* noise would similarly influence event-related neural responses during lexical processing. Specifically, we tested the hypothesis that age and neural network dynamics affect anticipatory processing.

### Neural Noise and Predictive Processing

Current approaches in cognitive neuroscience emphasize the role of active, top-down mechanisms in the processing of incoming sensory stimuli (e.g., Clark, 2013; Friston, 2005; Hinton, 2010). These models suggest that the brain proactively generates expectations for upcoming information, compares real-world input with these internally held expectations, and adjusts accordingly. Such generative models of anticipatory processing – or *prediction* – are thought to have vast explanatory power across cognitive domains, and therefore have become an increasingly popular assumption of computational networks modeling neural dynamics (e.g., Dayan, Hinton, Neal, & Zemel, 1995; Jaeger & Haas, 2004; Maloney & Mamassian, 2009). However, very little evidence exists to link prediction to specific neural activation patterns.

Emerging frameworks for anticipatory processing have described the brain as a Bayesian “prediction machine”, or a network that distills statistical regularities from environmental stimuli and uses these statistics to make predictions about future events (Bastos et al., 2012; Clark, 2013; Friston, 2010). The predictive brain probabilistically generates models for upcoming information, and is believed to pre-activate those models when information is highly expected (i.e., when the environment strongly supports presentation of a specific stimulus). In order for prediction to be both efficient and generalizable across sensory and cognitive domains (i.e., unifying; Friston, 2005), top-down processing must be a fundamental architectural feature of neural organization. One explanation for neural encoding of prediction suggests that anticipatory mechanisms are embedded in spatio-temporal relationships across neuronal populations (Engel, Fries, & Singer, 2001). This temporal binding model postulates that higher-order brain dynamics modulate synchronous timing of neural firing. In other words, top-down information (e.g., expectations, goals, and attention) influences the size, strength, and cohesive firing of neural assemblies activated for expected input. If so, synchronous firing should be enhanced during prediction. Recent evidence shows that synchronous oscillatory activity in the brain is associated with the presentation of predictable, as opposed to unpredictable, stimuli (reviewed in Engel, Fries, & Singer, 2001; Arnal, Wyart, & Giraud, 2011; Doelling, Arnal, Ghitza, & Poeppel, 2014; Samaha, Bauer, Cimaroli, & Postle, 2015). In order for networks to show neural synchrony, stable and coherent networks must be established in the brain (Fries, 2005), and so we propose a corollary: prediction is enhanced for individuals with more synchronous neuronal networks.

In the present study, we test this hypothesis by examining the effects of age and 1/*f* neural noise on ERP indices of predictive processing during language comprehension.

### Effects of Age and Noise on Prediction during Language Comprehension

ERP studies of lexical prediction typically examine effects of manipulating *cloze probability*, or the likelihood that a word will complete a given context^2^ (reviewed in Kutas & Federmeier, 2011). Effects of cloze probability have primarily been reported to modulate the amplitude of two ERP components: the N400 and, more recently, the post-N400 positivity (PNP). The N400 is a negative deflection that is maximal over posterior electrode sites between 300 to 500ms after a word is presented. The amplitude of the N400 is reduced when comprehenders process high-cloze (highly predictable) relative to low-cloze (unpredictable) words in sentence or discourse contexts (reviewed in Swaab, Ledoux, Camblin, & Boudewyn, 2012). The N400 effect is thought to index neural facilitation when words are highly expected or pre-activated. The PNP is maximal to low-cloze continuations over frontal electrode sites between 600-900ms (reviewed in Delong & Kutas, 2016; Van Petten & Luka, 2012). The PNP is thought to reflect costs of resolving an unexpected word with the prior discourse, or costs associated with updating the discourse representation with new information. Modulations of the N400 and PNP as a function of critical word predictability are found both as a function of if critical words were actually accurately predicted by the reader, or were plausible given the preceding context. An ERP paradigm developed by Brothers and colleagues (2015) allows us to observe separate N400 and PNP effects for prediction accuracy and contextual plausibility (independent of prediction accuracy, henceforth referred to as effects of *context*).

In the Brothers et al. study, participants were asked to try to predict the last word of moderate cloze (40–60%) two-sentence passages and, at the end of each passage, to report whether the passage-final word matched the identity of the word they had predicted. The recorded EEG was sorted in different bins for accurately predicted and inaccurately guessed words. A subset of the passages presented in this study ended in low-cloze (0–7%) completions, and readers nearly always reported inaccurately guessing these unpredictable items. By comparing ERP in three conditions (accurately predicted moderate-cloze, inaccurately predicted moderate-cloze, and inaccurately predicted low-cloze items), Brothers et al. (2015; see also Brothers et al., 2017) found separable N400 and PNP effects of accurate prediction and contextual support (i.e., cloze probability independent of prediction accuracy). These components were larger in amplitude when the reader’s prediction was incorrect and when the critical word was incongruent given the preceding context. Using this paradigm, Dave et al. replicated this pattern in older adult readers (Dave et al., a/b, submitted). Dave et al. further showed older adults had reduced effects of prediction accuracy and contextual support on the N400, while PNP effects were not modulated as a function of age. In the current study, we correlated ERP data collected in Dave et al. (b) with EEG measures of 1/*f* neural noise for each participant. Specifically, we examined whether separate ERP measures of prediction accuracy and contextual support correlated with individual estimates of 1/*f* neural noise.

## Methods

### Participants

#### Comprehension Paradigm

EEG data were collected from 24 young adults (19 females; mean age: 19.5; range: 18 to 28) and 24 older adults (14 females; mean age: 72.0 years; range 64 to 79). All participants were native English speakers with normal or corrected-to-normal vision and no known history of psychiatric or neurological disorders, head trauma, or neuro-active prescription medication. All participants were right-handed, as assessed by self-report and the Edinburgh Handedness Inventory (Oldfield, 1971). Both groups provided written informed consent to a protocol approved by the Institutional Review Board at the University of California, Davis. Data from two additional participants in each age group were excluded from analyses due to excessive artefacts in EEG recordings.

#### Prediction Paradigm

EEG data were collected from 36 new young adult participants (19 females, mean age: 20.5; range: 18 to 33). The same 24 older adults were tested on the prediction paradigm, and 12 new older adult participants were also included (total group: 21 females; mean age: 70.8 years; range 64 to 79). All participants were consented to the same criteria described above. Data from two additional young adults and three additional older adults were excluded from the analyses as a result of excessive artefacts in EEG recordings.

### Stimuli and Procedure

#### Comprehension Paradigm

Participants read 100 experimental sentences, presented via rapid serial visual presentation (RSVP) with a stimulus-onset asynchrony (SOA) of 600ms (as described in Dave et al., a, submitted). Participants were instructed to read for comprehension, which was tested by manual response to true/false statements following 25 percent of the trials.

#### Prediction Paradigm

Participants read 180 two-sentence discourse passages (as described in Brothers et al., 2015). Sample stimuli for the Prediction Paradigm are presented in Table 1.

**Table 1.** Sample experimental stimuli for the Prediction Paradigm.

| **Prediction Task Paradigm** |  |  |  |  |
| --- | --- | --- | --- | --- |
| *Condition* | *Sentence 1* | *Sentence 2* | *Expectation* | *Final Word* |
| *Moderate Cloze, Accurately Predicted* | We wanted a traditional pet in the apartment. | That's why we bought a... | Dog | <u>Dog</u> |
| *Moderate Cloze, Inaccurately Predicted* | We wanted a traditional pet in the apartment. | That's why we bought a... | Cat | <u>Dog</u> |
| *Low Cloze, Inaccurately Predicted* | Our family needed to protect our diamonds and jewelry. | That's why we bought a... | Safe | <u>Dog</u> |

The first sentence of each passage was presented all at once, and the second sentence was presented via RSVP (SOA: 600ms). The materials consisted of 120 passages ending with a moderate cloze target word (40–60%, average: 50.7%), and 60 passages ending with a low cloze target word (0–7%, average: 0.9%). Participants were instructed to try to use the discourse context to predict the passage final word. Readers were informed that there were no “correct” predictions for any passage. After each passage, readers manually indicated whether the actually presented passage-final word matched their prediction.

### EEG Recording

EEG data was recorded using SCAN (Compumedics Neuroscan), and processing and analysis was performed using Matlab (The MathWorks, Natick, MA), with the EEGLAB toolbox. EEG was recorded during both tasks with 29 tin electrodes mounted in an elastic cap (ElectroCap International). Additional electrodes were attached below and on the outer canthi of the eyes in order to capture blinks and other eye movements. Electrode impedances were kept below 5kΩ, and the EEG signal was amplified with a Synamps Model 8050 Amplifier (bandpass cutoffs: 0.05 – 100 Hz). The signal was continuously digitized at a sampling rate of 250 Hz.

### EEG Analysis for *1*/f Neural Noise

For data collected during both Comprehension and Prediction Paradigms, EEG was epoched to exclude all time during which participants were not on task (e.g., rest or breaks between sentences). We replicated the broadband power spectral density (PSD) analyses for 1/*f* neural noise originally performed by Voytek et al. (2015). Voytek and colleagues estimated slope from EEG in the 2-25 Hz PSD range, excluding the alpha range (7-14 Hz). High frequency activity (gamma band; 25-100 Hz) was excluded from scalp recordings due to likely overlap with face and eye muscle activity (e.g., Voytek et al., 2010; Luck, 2005). Alpha activity was excluded from slope analyses because oscillations in this frequency band are inconsistent with broadband local-field potential patterns (i.e., alpha activity is representative of non-broadband activity; Manning et al., 2009; Miller, 2010; Miller et al., 2014).

As in Voytek et al. (2015), we extracted PSD at each electrode site for each participant. PSD is thought to be inversely proportional with frequency (Miller, 2010), and therefore PSD was log transformed. By using log_10_PSD, linear regressions generated between frequency and PSD yielded a representative slope (i.e., 1/*f* neural noise slope).

### ERP Analysis for the Prediction Paradigm

ERP processing was performed using Matlab with the EEGLAB toolbox and ERPlab plugin (Luck, 2005). After EEG recording, independent components analysis (ICA) was used to decompose EEG responses into subcomponents with fixed scalp distributions and independent time courses. After ICA, eye blink components were removed, and single-trial waveforms were screened for amplifier blocking, muscle artifacts, and horizontal eye movements.

ERP waveforms were first sorted into three conditions: (i) accurately predicted moderate cloze completions, (ii) inaccurately predicted moderate cloze completions, and (iii) inaccurately predicted low cloze completions. These waveforms are plotted in Figure 1A. Difference waveforms were plotted (Figure 1B) to display two effects: Prediction (unpredicted minus predicted moderate cloze targets) and Context (unpredicted low minus unpredicted moderate cloze targets).

**Figure 1.**
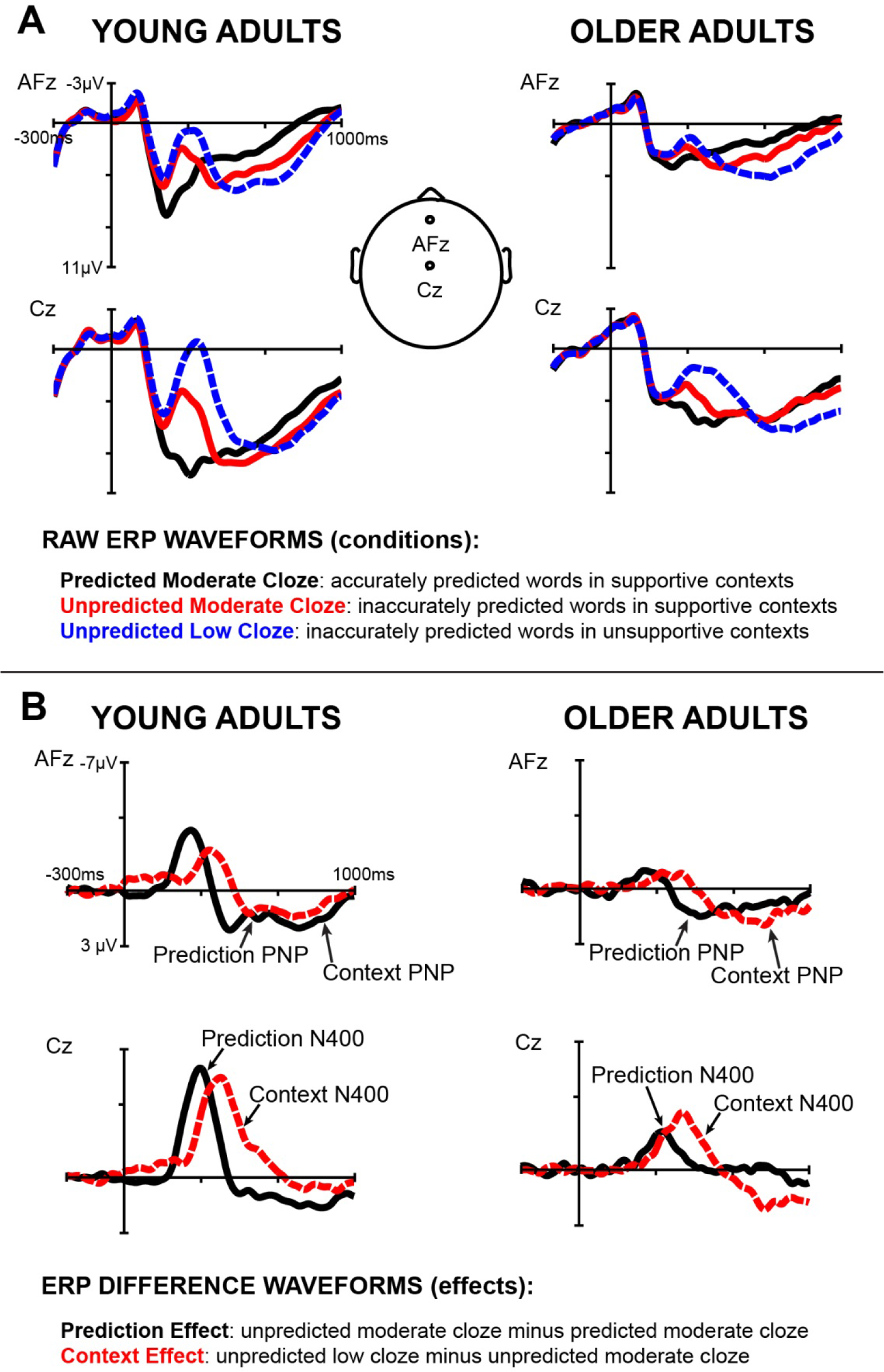
ERP results. (**A**) Averaged event-related potentials for passage-final critical words are displayed for young (left) and older adult (right) readers in the Prediction Paradigm. (**B**) Difference waveforms were generated by subtracting between ERP conditions to display effects of Prediction and Context.

As seen in Brothers et al. (2015; 2017) and Dave et al. (a/b, submitted), young and older adults showed separate negative deflections between 300-500ms for effects of Prediction and Context. These effects were maximal at the typical centro-parietal sites (i.e., reviewed in Kutas & Federmeier, 2011; Swaab, Ledoux, Camblin, & Boudewyn, 2012). Both age groups also showed significant positive deflections between 600-900ms for both the effects of Prediction and Context. This post-N400 positivity (PNP) was maximal over frontal electrode sites (Brothers et al., 2015; 2017; Federmeier, Wlotko, Ochoa-Dewald & Kutas, 2007; Van Petten & Luka, 2012).

Older adults typically show delayed ERP effects relative to younger readers (meta-analysis in Kutas & Iragui, 1998), and in the present study we also observed delayed N400 and PNP effects in older relative to young adults in the Prediction Paradigm (Figure 1B). Therefore, as described in Dave et al. (a/b, submitted), ERP effects were measured over 100ms epochs on maximal peak amplitudes for each group. N400 effects were measured over a cluster of six centro-posterior electrode sites (CP1/2, P3/4, Cz, Pz), while PNP effects were calculated across a cluster of six frontal electrode sites (FP1/2, F3/4, AFz, Fz)

## Results

### *1*/f Neural Noise across Age in Comprehension and Prediction Paradigms

We generated linear regressions modeling the relationship between frequency and PSD (log_10_PSD), and analyzed the slopes (1/*f* neural noise) emerging from these analyses. Significant differences in 1/*f* neural noise were observed between older and young adults on both comprehension and prediction tasks (Figure 2A). In both tasks, younger adults had significantly steeper slopes than older adults (comprehension: *r*(47) = .613, p < .001; prediction: *r*(47) = .483, p < .001; replicating the result of similar analyses in Voytek et al., 2015). Regression analyses provided good fit of the data in both age groups and tasks (*r*s > .70), indicating linear modeling of the data is well-motivated.

**Figure 2.**
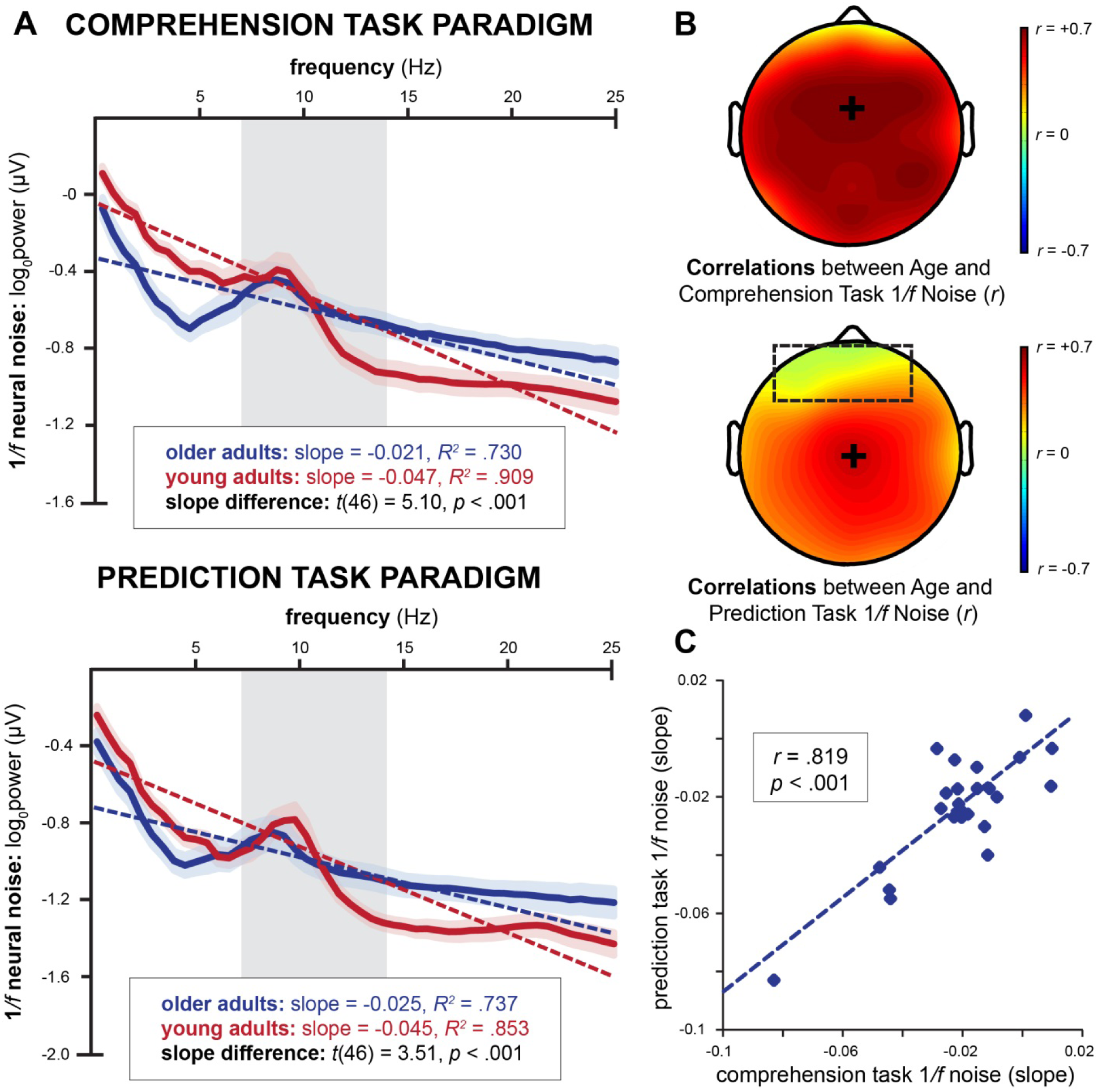
1/*f* neural noise results. (**A**) 1/*f* neural noise was estimated for both Comprehension and Prediction Paradigms from the slope (dotted lines) of the power spectrum across frequency (2-25 Hz, plotted above with 95% confidence intervals), excluding alpha frequency (7-14 Hz, shaded). Correlations between Age and 1/*f* slopes are plotted topographically (**B**). Correlations were maximal over central electrode sites for both paradigms (addition signs). Age correlations with neural noise were not significantly different as a function of task, but topographic differences between tasks emerged over frontal electrode sites (dotted outline). (**C**) 1/*f* slopes were strongly correlated between tasks for older readers.

In order to address whether 1/*f* neural noise slopes were similar between tasks, we first examined the frequencies for which age-related differences were significant. In the comprehension task, significant age differences emerged across the entire 2-25 Hz scale, excluding a subset of the alpha range (7-12.3 Hz, *p* range:.12 to.44). Power decreased with age at lower frequencies (2-7 Hz, high delta and theta), but increased with age at higher frequencies (beta). In contrast, significant age differences only emerged in a higher, primarily beta frequency range in the prediction task (13.3-25 Hz). As in the comprehension task, older adults had high power than younger adults at higher frequencies (*p*s > .01).

Next, we examined topographic correlations between age and 1/*f* noise, to observe where age-related differences in neural noise were most pronounced across the scalp (Figure 2B). In the comprehension task, age differences were maximal over much of the scalp, with the strongest age correlations emerging over central electrode sites. While correlations between age and 1/*f* noise were generally reduced for the prediction relative to the comprehension task, these differences were not significant (whole head: *Z* = 1.21, *p* = .23). As displayed in Figure 2B, task differences in noise-age correlations only emerged significantly over frontal electrode sites (comprehension task average: *r* = .598, prediction task average: *r* = .233, Z = 2.15, *p* = .03).

As 1/*f* slopes varied as a function of language task across both frequency and topography, we aimed to determine if participants showed intra-individual reliability in 1/*f* neural noise (i.e., if slope measured across the entire scalp correlated for both tasks). 24 older adults performed both comprehension and prediction tasks^3^, and served as our test population for evaluating whether 1/*f* neural noise was a reliable measure of individual difference. As displayed in Figure 2C, slopes were significantly correlated (*r* = .819, *p* < .001) between tasks for older adults, suggesting that 1/*f* neural noise remains stable despite variable task demands.

### Mediation Analyses for Age, *1*/f Neural Noise, and ERP Effects

We next assessed whether ERP effect amplitudes (displayed in Figure 1B) were impacted by 1/*f* slopes, or Noise averaged across the scalp. In order to do so, we first correlated ERP effect amplitudes with continuous variables of Age and Noise, and then performed mediation analyses (Sobel, 1982) to examine whether Noise mediated Age effects on ERP amplitudes. N400 effects of both Prediction and Context were significantly reduced in older readers (*p*s < .001, Figure 3A), while PNP effects were not impacted by Age (*p*s > .5). Likewise, flatter 1/*f* slopes correlated with smaller N400 effects of both Prediction (*p* < .001) and Context (*p* = .007, Figure 3B), while no significant correlations were found for PNP effects. As both Age and Noise were predictive of N400 amplitudes, and Age correlated significantly with Noise (*R^2^* = .240, *F*(1, 70) = 22.09, *p* < .001), independent models were constructed to examine whether Noise mediated effects of Age on N400 effects of Prediction or Context (Figure 3C).

**Figure 3.**
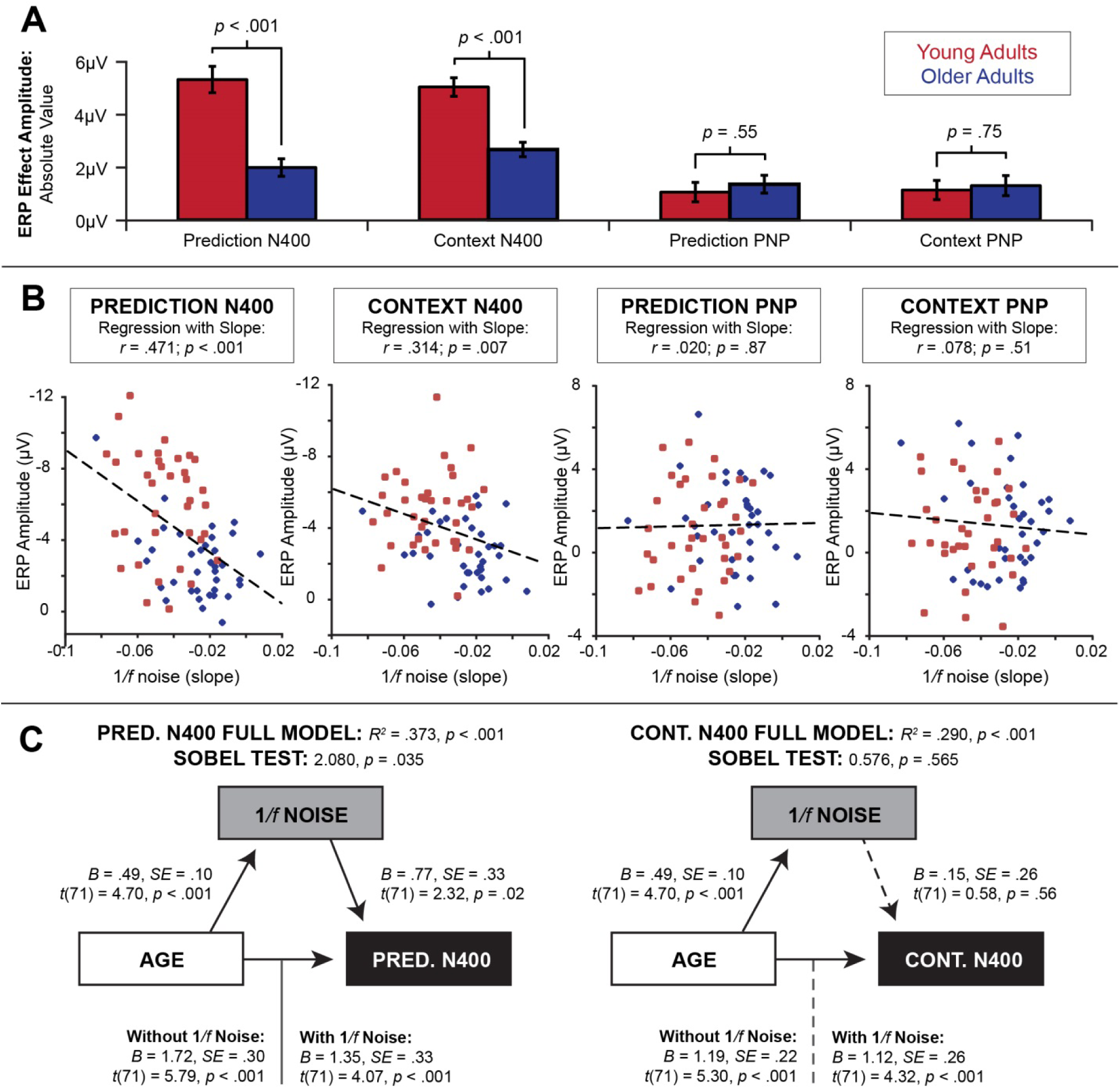
Age, 1/*f* noise, and ERP effects. N400 amplitudes for Prediction and Context were reduced in older readers (blue) relative to younger adults (red, **A**), but no significant effects of age were found for either of the PNP effects. (Error bars indicate standard errors (SEM).) 1/*f* neural noise was significantly correlated with amplitudes of both N400 effects (**B**), but not with the PNP effects. A mediation model generated for the Prediction N400 (**C**, left) shows that 1/*f* neural noise partially mediated age-related reductions in the Prediction N400. In contrast, a similar mediation analysis performed for the Context N400 (**C**, right) shows that 1/*f* neural noise is not predictive of Context N400 amplitudes after controlling for age, and does not mediate age-related reduction of the Context N400 (dotted lines).

Age significantly predicted the N400 effect of Prediction (*t*(71) = 5.79, *p* < .001) in a model generated without Noise, *R^2^* = .324, *F*(1, 70) = 33.56, *p* < .001. In a multiple regression analysis with both Age and Noise, the effect of Age on Prediction N400 remained significant (*t*(71) = 4.07, *p* < .001) but was partially mediated by Noise (Sobel test statistic: 2.08, *p* = .035). In the presence of Age, Noise still significantly predicted Prediction N400, *t*(71) = 2.32, *p* = .02. While Age alone explained 32% of the total variance in Prediction N400 amplitude, an additional 5% of the variance was explained by effects of Noise unrelated to Age, and this addition to model fit was significant.

Similar analyses were performed for effects of Age and Noise on Context N400. Age significantly predicted the N400 effect of Prediction (*t*(71) = 5.30, *p* < .001) in a model generated without Noise, *R^2^* = .287, *F*(1, 70) = 28.12, *p* < .001. With the addition of Noise to the model, the effect of Age on Prediction N400 remained significant (*t*(71) = 4.32, *p* < .001) and Noise was not found to mediate this relationship (Sobel test statistic: 0.58, *p* = .565). Noise was not predictive of Context N400 in the presence of Age, *t*(71) = 0.58, *p* = .56, and only 0.3% additional variance was explained by effects of Noise unrelated to Age in this model.

In addition to mediation models, we conducted two additional sets of analyses. As observed above, age-related 1/*f* slope differences in the Prediction Paradigm were primarily driven by increased power in the beta range (14-25 Hz) for older adults. We therefore examined whether age-related differences in beta activity correlated with N400 amplitudes, to assess if 1/*f* noise slope correlations with N400 amplitudes were driven by beta power. However, average power in the beta band was not significantly correlated with either prediction accuracy (*r* = .038, *p* = .10) or contextual support (*r* = .007, *p* = .47) effects on the N400.

Secondly, we investigated if Age significantly moderated the effect of Noise on N400 amplitudes, to test if different relationships between Noise and N400 effects emerged in each age group. However when entered alongside Age and Noise, the Noise by Age interaction was not significantly predictive of N400 effects of either Prediction (*t*(71) = −0.02, *p* = .99) or Context (*t*(71) = 0.84, *p* = .40).

## Discussion

Consistent with previous findings, the current study demonstrates that 1/*f* neural noise reliably changes as a function of normative aging. Across both studies, slopes of the 1/*f* function flattened with advanced age, showing that this marker can be observed in EEG recorded in tasks with different cognitive demands. Prior computational models (Freeman & Zhai, 2009; Manning et al., 2009; Miller et al., 2007) have suggested that this broadband measure of 1/*f* noise may reflect the degree to which neural networks are synchronized in their firing, proposing that observed 1/*f* noise differences may reflect neuronal desynchronization in aging (i.e., neural noise hypothesis, Crossman & Szafram, 1956).

While young adults typically had more negative noise slopes than older adults, slope ranges showed some overlap between groups. Therefore, while neural noise correlated with age, 1/*f* slopes were not entirely explained by age group differences. We observed age effects on noise slopes in each task, and found two key results. First, age correlated most strongly with 1/*f* slopes in central regions of the scalp for both tasks (as in Onton, Delorme, & Makeig, 2005; Rajah & D’Esposito, 2005). Interestingly, the addition of the prediction task resulted in a shifted topography for age by slopes correlations, suggesting that age differences in neural activation patterns are modulated by additional task demands (as seen in Gazzaley et al., 2008; Klostermann et al., 2007). Second, we correlated noise slopes across language tasks in the population of older adults studied in the current experiment. Whole head 1/*f* slopes strongly correlated despite both different reading goals and different topographies between tasks, reflecting that noise is a consistent and characteristic measure of the individual. These results imply that 1/*f* noise may serve as a neural measure of individual differences in older cohorts. Between-task correlations in noise for young adults could not be calculated in the present study, as two groups of young adults participated in the two experiments. It will be important for future studies to examine whether similar 1/*f* noise correlations emerge in both age groups.

Expanding on the findings of Voytek and colleagues (2015), our study aimed to investigate whether the steepness of the 1/*f* slopes was correlated with ERP measures related to language processing. Our results show that N400 effects were modulated by this EEG-derived measure of noise, but PNP effects were not correlated with noise. Therefore, the effects of neural noise may be specific to certain language comprehension processes. N400 modulations are associated with facilitation of lexical-semantic processing when a presented word is accurately predicted (Prediction N400), or matches the meaning of the previous context independent of prediction accuracy (Context N400). In contrast, PNP amplitudes have been associated with “work” necessary to revise discourse representations (Brothers et al., 2015; Van Petten & Luka, 2012) following the presentation of unexpected words. Our findings suggest that these updating processes are less likely to be affected by neural synchrony.

A key finding of the current study is that 1/*f* neural noise specifically influenced N400 effects of accurate lexical prediction. This result is largely consistent with Engel et al.’s (2010) proposal that neural network dynamics – specifically, population spiking synchrony – underlie the strength of predictions generated during anticipatory processing. Several studies have found that oscillatory activity in narrowband frequency ranges change for predictable, as opposed to unpredictable, stimuli during language comprehension (Lewis & Bastiaansen, 2015; Rommers et al., 2016; Vignali et al., 2016). Critically, these studies have emphasized the “frequency [band] specificity” of these findings (Bastiaansen & Hagoort, 2016, p. 190). However, our results indicate that observed age-related differences in spectral power in specific frequency bands (i.e., beta) did not correlate with N400 effects of predictive accuracy, and therefore cannot explain how 1/*f* neural noise mediates age-related changes in this neural index of prediction. Therefore, our current study re-affirms Voytek et al. (2015)’s argument that researchers benefit from studying the power spectrum as a “unified statistical representation of the signal” (p. 13262).

To our knowledge, no prior studies have examined the role of broadband EEG activity in predictive processing. Furthermore, our study is the first to find individual differences in predictive processing as a function of neural noise and aging. This EEG-derived measure represents a readily available, non-invasive index of neural synchrony that can be observed at the single subject level to predict individual differences in behavioural and neural outcomes. There is broad potential for future studies in applying this technique in understanding individual differences in cognitive processes, both in typical populations and in individuals affected by neural disease (i.e., Alzheimer’s). However, it remains to be determined whether an accurate index of 1/*f* noise can be obtained over shorter time scales, or in resting state. Additional research will be necessary for determining the recording characteristics that will optimize 1/*f* neural noise measurement for laboratory and clinical investigation.

It is particularly important to note that in both Voytek et al. and the current study, 1/*f* slopes did not mediate all behavioral or neural measures showing age-related declines (e.g., no mediation was found for response times in Voytek et al., or N400 effects of Context in the current study). Instead, noise mediated specific age effects of (i) accurate behavioral responses and (ii) neural effects of prediction accuracy. Friston (2005)’s foundational model of predictive processing defines accuracy as a minimization of prediction error, yielding reductions in bottom-up neural activity when a comprehender has pre-activated (predicted) a representation of the expected stimulus. This conceptualization suggests that neural noise slopes may specifically reflect the maintenance of top-down mechanisms that allow comprehenders to generate expectations and/or pre-activate specific stimuli (as posited by Engel et al., 2010; Clark, 2013). It will be critical for future research to address how neural synchrony modulates top-down processing. Computational and theoretical models have suggested 1/*f* noise may dynamically influence effortful processing (Grigolini et al., 2009), representation formation (Gilden, 2001), and strategic shifting (reviewed in Wagenmakers, Farrell, & Ratcliff, 2004) – all of which have been suggested to underlie predictive processing across the lifespan (i.e., Clark, 2013; Federmeier, 2007; Kuperberg & Jaeger, 2016).

Because N400 effects of Context were not linked to 1/*f* neural noise after controlling for age, activation patterns required for effective contextual processing are very likely distinct from those underlying predictive mechanisms (as seen in oscillatory analyses performed by Rommers et al., 2016). Importantly, no evidence emerged to suggest differential involvement of 1/*f* noise in predictive processing in young and older adults. Instead, we posit that similar neural mechanisms for lexical prediction (Kuperberg & Jaeger, 2016) are recruited by both young and older adults, and that benefits of top-down information are guided by both neuronal spiking synchrony, as well as sources of variance that are related to age but may be independent of 1/*f* neural noise.

What these sources of variance may be is an open question, but neural noise as measured by EEG 1/*f* power alone may not fully describe how noise modulates neural network dynamics. Some researchers have posited that inter-trial variability may also capture age-related decline in signal fidelity, and have successfully linked aging to inconsistency in behavioral responses (Hong & Rebec, 2012; Hultsch & MacDonald, 2004; Lovden, Li, Shing, & Lindenberger, 2007), as well as fMRI (D’Esposito et al., 1999) and oscillatory phase coherence (Papenberg et al., 2013). Further, recent evidence (Payne & Federmeier, 2017) has emerged to suggest that contextual processing may be dependent on intra-individual, trial-to-trial variability in ERP activity. Taken together with the results of the current study, Payne and Federmeier’s results indicate that effects of prediction accuracy and contextual support may both be modulated noise, but that these measures represent distinct subcomponents of neural noise. Future research would benefit from incorporating elements of both network-level spiking synchronization (i.e., 1/*f* neural noise) and trial-specific neural activity (i.e., inter-trial variability) in gauging the mechanisms underlying language processing in young and older adults.

## Footnotes

1 1/*f* noise, alternately known as *flicker* or *pink nose*, is defined as electrical activity associated with current flow in systems with few charge carriers, such as neurons (e.g., Hooge & Gaal, 1971; Stevens, 1972). 1/*f* neural noise has been observed in mammalian neural circuits since Verveen and Derksen (1965).

2 For example, for the sentence: *You could tell he was angry from the tone of his*…, the most predictable and therefore highest cloze completion is *voice*. An example of an unpredictable, low cloze competition is *violin.*

3 Correlations between tasks were not calculated for young adults because two separate sets of young readers participated in the Comprehension and Prediction Paradigm tasks.

## References

Arnal, L. H., Wyart, V., & Giraud, A. L. (2011). Transitions in neural oscillations reflect prediction errors generated in audiovisual speech. Nature Neuroscience, 14(6), 797–801.

Bastiaansen, M., & Hagoort, P. (2006). Oscillatory neuronal dynamics during language comprehension. Progress in Brain Research, 159, 179–196.

Bastos, A. M., Usrey, W. M., Adams, R. A., Mangun, G. R., Fries, P., & Friston, K. J. (2012). Canonical microcircuits for predictive coding. Neuron, 76(4), 695–711.

Brothers, T., Swaab, T. Y., & Traxler, M. J. (2015). Effects of prediction and contextual support on Lexical processing: Prediction takes precedence. Cognition, 136, 135–149.

Brothers, T., Swaab, T. Y., & Traxler, M. J. (2017). Goals and strategies influence lexical prediction during sentence comprehension. Journal of Memory and Language, 93, 203–216.

Cabeza, R., Nyberg, L., & Park, D. C. (Eds.). (2016). Cognitive neuroscience of aging: Linking cognitive and cerebral aging. Oxford University Press.

Clark, A. (2013). Whatever next? Predictive brains, situated agents, and the future of cognitive science. Behavioral and Brain Sciences, 36(03), 181–204.

Cremer, R., & Zeef, E. J. (1987). What kind of noise increases with age?. Journal of Gerontology, 42(5), 515–518.

Crossman, E. R., & Szafran, J. (1955). Changes with age in the speed of information-intake and discrimination. Experientia, 4, 128–34.

D’Esposito, M., Postle, B. R., Jonides, J., & Smith, E. E. (1999). The neural substrate and temporal dynamics of interference effects in working memory as revealed by event-related functional MRI. Proceedings of the National Academy of Sciences, 96(13), 7514–7519.

Dave, S., Brothers, T.A., Traxler, M.J., Ferreira, F., Henderson, J.M., & Swaab, T.Y. (a, submitted). Electrophysiological evidence for preservation of lexical prediction in aging.

Dave, S., Brothers, T.A., Long, D., & Swaab, T.Y. (b, submitted). Individual differences in lexical prediction across the lifespan: Effects of verbal fluency, vocabulary skill and working memory.

Dayan, P., Hinton, G. E., Neal, R. M., & Zemel, R. S. (1995). The Helmholtz machine. Neural Computation, 7(5), 889–904.

DeLong, K. A., & Kutas, M. (2016). Hemispheric differences and similarities in comprehending more and less predictable sentences. Neuropsychologia, 91, 380–393.

DeLong, K. A., Groppe, D. M., Urbach, T. P., & Kutas, M. (2012). Thinking ahead or not? Natural aging and anticipation during reading. Brain and Language, 121(3), 226–239.

Doelling, K. B., Arnal, L. H., Ghitza, O., & Poeppel, D. (2014). Acoustic landmarks drive delta– theta oscillations to enable speech comprehension by facilitating perceptual parsing, Neuroimage, 85, 761–768.

Engel, A. K., Fries, P., & Singer, W. (2001). Dynamic predictions: oscillations and synchrony in top–down processing. Nature Reviews Neuroscience, 2(10), 704–716.

Federmeier, K. D. (2007). Thinking ahead: The role and roots of prediction in language comprehension. Psychophysiology, 44(4), 491–505.

Federmeier, K. D., Kutas, M., & Schul, R. (2010). Age-related and individual differences in the use of prediction during language comprehension. Brain and Language, 115(3), 149–161.

Federmeier, K. D., McLennan, D. B., Ochoa, E., & Kutas, M. (2002). The impact of semantic memory organization and sentence context information on spoken language processing by younger and older adults: An ERP study. Psychophysiology, 39(2), 133–146.

Federmeier, K. D., Wlotko, E. W., De Ochoa-Dewald, E., & Kutas, M. (2007). Multiple effects of sentential constraint on word processing. Brain Research, 1146, 75–84.

Fjell, A. M., & Walhovd, K. B. (2010). Structural brain changes in aging: courses, causes and cognitive consequences. Reviews in the Neurosciences, 21(3), 187–222.

Freeman, W. J., & Zhai, J. (2009). Simulated power spectral density (PSD) of background electrocorticogram (ECoG). Cognitive Neurodynamics, 3(1), 97–103.

Fries, P., Nikolić, D., & Singer, W. (2007). The gamma cycle. Trends in Neurosciences, 30(7), 309–316.

Fries, P. (2005). A mechanism for cognitive dynamics: neuronal communication through neuronal coherence. Trends in Cognitive Sciences, 9(10), 474–480.

Friston, K. (2005). A theory of cortical responses. Philosophical Transactions of the Royal Society of London B: Biological Sciences, 360(1456), 815–836.

Friston, K. (2010). The free-energy principle: a unified brain theory?. Nature Reviews Neuroscience, 11(2), 127–138.

Gazzaley, A., Clapp, W., Kelley, J., McEvoy, K., Knight, R. T., & D’Esposito, M. (2008). Age-related top-down suppression deficit in the early stages of cortical visual memory processing. Proceedings of the National Academy of Sciences, 105(35), 13122–13126.

Gilden, D. L. (2001). Cognitive emissions of 1/*f* noise. Psychological Review, 108, 33–56.

Grigolini, P., Aquino, G., Bologna, M., Luković, M., & West, B. J. (2009). A theory of 1/*f* noise in human cognition. Physica A: Statistical Mechanics and its Applications, 388(19), 4192–4204.

He, B. J. (2014). Scale-free brain activity: past, present, and future. Trends in cognitive sciences, 18(9), 480–487.

Hinton, G. (2010). A practical guide to training restricted Boltzmann machines. Momentum, 9(1), 926–946.

Hong, S. L., & Rebec, G. V. (2012). A new perspective on behavioral inconsistency and neural noise in aging: compensatory speeding of neural communication. Frontiers in Aging Neuroscience, 4, 27.

Hooge, F. N., & Gaal, J. L. M. (1971). Fluctuations with a 1/*f* spectrum in conductance of ionic solutions and in voltage of concentration cells. Philips Research Reports, 26(2), 77.

Huettig, F., & Janse, E. (2016). Individual differences in working memory and processing speed predict anticipatory spoken language processing in the visual world. Language, Cognition and Neuroscience, 31(1), 80–93.

Hultsch, D. F., & MacDonald, S. W. (2004). Intraindividual variability in performance as a theoretical window onto cognitive aging. *New Frontiers in Cognitive Aging*, 65–88.

Jaeger, H., & Haas, H. (2004). Harnessing nonlinearity: Predicting chaotic systems and saving energy in wireless communication. Science, 304(5667), 78–80.

Klostermann, F., Nikulin, V. V., Kühn, A. A., Marzinzik, F., Wahl, M., Pogosyan, A., & Curio, G. (2007). Task‐related differential dynamics of EEG alpha‐and beta‐band synchronization in cortico‐basal motor structures. European Journal of Neuroscience, 25(5), 1604–1615.

Kreiman, G., Hung, C. P., Kraskov, A., Quiroga, R. Q., Poggio, T., & DiCarlo, J. J. (2006). Object selectivity of local field potentials and spikes in the macaque inferior temporal cortex. Neuron, 49(3), 433–445.

Kuperberg, G. R., & Jaeger, T. F. (2016). What do we mean by prediction in language comprehension?. Language, Cognition and Neuroscience, 31(1), 32–59.

Kutas, M., & Federmeier, K. D. (2011). Thirty years and counting: finding meaning in the N400 component of the event-related brain potential (ERP). Annual Review of Psychology, 62, 621–647.

Kutas, M., & Iragui, V. (1998). The N400 in a semantic categorization task across 6 decades. Electroencephalography and Clinical Neurophysiology/Evoked Potentials Section, 108(5), 456–471.

Lewis, A. G., & Bastiaansen, M. (2015). A predictive coding framework for rapid neural dynamics during sentence-level language comprehension. Cortex, 68, 155–168.

Li, S. C., Lindenberger, U., & Sikström, S. (2001). Aging cognition: from neuromodulation to representation. Trends in Cognitive Sciences, 5(11), 479–486.

Lövdén, M., Li, S. C., Shing, Y. L., & Lindenberger, U. (2007). Within-person trial-to-trial variability precedes and predicts cognitive decline in old and very old age: Longitudinal data from the Berlin Aging Study. Neuropsychologia, 45(12), 2827–2838.

Luck, S. J. (2005). Ten simple rules for designing ERP experiments. *Event-related potentials: A methods handbook*, 262083337.

Maloney, L. T., & Mamassian, P. (2009). Bayesian decision theory as a model of human visual perception: testing Bayesian transfer. Visual Neuroscience, 26(01), 147–155.

Manning, J. R., Jacobs, J., Fried, I., & Kahana, M. J. (2009). Broadband shifts in local field potential power spectra are correlated with single-neuron spiking in humans. Journal of Neuroscience, 29(43), 13613–13620.

Miller, K. J. (2010). Broadband spectral change: evidence for a macroscale correlate of population firing rate?. Journal of Neuroscience, 30(19), 6477–6479.

Miller, K. J., Honey, C. J., Hermes, D., Rao, R. P., & Ojemann, J. G. (2014). Broadband changes in the cortical surface potential track activation of functionally diverse neuronal populations. Neuroimage, 85, 711–720.

Miller, K. J., Leuthardt, E. C., Schalk, G., Rao, R. P., Anderson, N. R., Moran, D. W., & Ojemann, J. G. (2007). Spectral changes in cortical surface potentials during motor movement. Journal of Neuroscience, 27(9), 2424–2432.

Nevin, J. A. (1969). Signal Detection Theory and Operant Behavior: A Review of David M. Green and John A. Swets’ Signal Detection Theory and Psychophysics. Journal of the Experimental Analysis of Behavior, 12(3), 475–480.

Oldfield, R. C. (1971). The assessment and analysis of handedness: the Edinburgh inventory. Neuropsychologia, 9(1), 97–113.

Onton, J., Delorme, A., & Makeig, S. (2005). Frontal midline EEG dynamics during working memory. Neuroimage, 27(2), 341–356.

Papenberg, G., Hämmerer, D., Müller, V., Lindenberger, U., & Li, S. C. (2013). Lower theta inter-trial phase coherence during performance monitoring is related to higher reaction time variability: a lifespan study. NeuroImage, 83, 912–920.

Payne, B. R., & Federmeier, K. D. (2017). Pace yourself: Intraindividual variability in context use revealed by self-paced event-related brain potentials. Journal of Cognitive Neuroscience, 29(5), 837–854.

Peelle, J. E., Troiani, V., Wingfield, A., & Grossman, M. (2010). Neural processing during older adults’ comprehension of spoken sentences: age differences in resource allocation and connectivity. Cerebral Cortex, 20(4), 773–782.

Podvalny, E., Noy, N., Harel, M., Bickel, S., Chechik, G., Schroeder, C. E., & Malach, R. (2015). A unifying principle underlying the extracellular field potential spectral responses in the human cortex. Journal of Neurophysiology, 114(1), 505–519.

Rajah, M. N., & D’esposito, M. (2005). Region-specific changes in prefrontal function with age: a review of PET and fMRI studies on working and episodic memory. Brain, 128(9), 1964–1983.

Raz, N., Lindenberger, U., Rodrigue, K. M., Kennedy, K. M., Head, D., Williamson, A., & Acker, J. D. (2005). Regional brain changes in aging healthy adults: general trends, individual differences and modifiers. Cerebral Cortex, 15(11), 1676–1689.

Rommers, J., Dickson, D. S., Norton, J. J., Wlotko, E. W., & Federmeier, K. D. (2017). Alpha and theta band dynamics related to sentential constraint and word expectancy. Language, Cognition and Neuroscience, 32(5), 576–589.

Salthouse, T. A., & Lichty, W. (1985). Tests of the neural noise hypothesis of age-related cognitive change. Journal of Gerontology, 40(4), 443–450.

Samaha, J., Bauer, P., Cimaroli, S., & Postle, B. R. (2015). Top-down control of the phase of alpha-band oscillations as a mechanism for temporal prediction. Proceedings of the National Academy of Sciences, 112(27), 8439–8444.

Serletis, D., Zalay, O. C., Valiante, T. A., Bardakjian, B. L., & Carlen, P. L. (2011). Complexity in neuronal noise depends on network interconnectivity. Annals of Biomedical Engineering, 39(6), 1768–1778.

Sobel, M. E. (1982). Asymptotic confidence intervals for indirect effects in structural equations models. In S. Leinhart (Ed.), Sociological Methodology (pp. 290–312).

Stevens, C. F. (1972). Inferences about membrane properties from electrical noise measurements. Biophysical Journal, 12(8), 1028–1047.

Swaab, T. Y., Ledoux, K., Camblin, C. C., & Boudewyn, M. A. (2012). Language-related ERP components. Oxford Handbook of Event-Related Potential Components, 397–440.

Swets, J. A. (2014). Signal detection theory and ROC analysis in psychology and diagnostics: Collected papers. Psychology Press.

Wagenmakers, E. J., Farrell, S., & Ratcliff, R. (2004). Estimation and interpretation of 1/*f^α^* noise in human cognition. Psychonomic Bulletin & Review, 11(4), 579–615.

Van Petten, C., & Luka, B. J. (2012). Prediction during language comprehension: Benefits, costs, and ERP components. International Journal of Psychophysiology, 83(2), 176–190.

Verveen, A. A., & Derksen, H. E. (1965). Fluctuations in membrane potential of axons and the problem of coding. Biological Cybernetics, 2(4), 152–160.

Vignali, L., Himmelstoss, N. A., Hawelka, S., Richlan, F., & Hutzler, F. (2016). Oscillatory brain dynamics during sentence reading: A Fixation-related spectral perturbation analysis. *Frontiers in Human Neuroscience*, 10.

Voytek, B., Canolty, R. T., Shestyuk, A., Crone, N., Parvizi, J., & Knight, R. T. (2010). Shifts in gamma phase–amplitude coupling frequency from theta to alpha over posterior cortex during visual tasks. Frontiers in Human Neuroscience, 4, 191.

Voytek, B., & Knight, R. T. (2015). Dynamic network communication as a unifying neural basis for cognition, development, aging, and disease. Biological Psychiatry, 77(12), 1089–1097.

Voytek, B., Kramer, M. A., Case, J., Lepage, K. Q., Tempesta, Z. R., Knight, R. T., & Gazzaley, A. (2015). Age-related changes in 1/*f* neural electrophysiological noise. Journal of Neuroscience, 35(38), 13257–13265.

Welford, A. T. (1981). Signal, noise, performance, and age. Human Factors, 23(1), 97–109.

Wlotko, E. W., Federmeier, K. D., & Kutas, M. (2012). To predict or not to predict: Age-related differences in the use of sentential context. Psychology and Aging, 27(4), 975–988.

